# Decoding the orientation of contrast edges from MEG evoked and induced responses

**DOI:** 10.1101/148056

**Authors:** Dimitrios Pantazis, Mingtong Fang, Sheng Qin, Yalda Mohsenzadeh, Quanzheng Li, Radoslaw Martin Cichy

**Author notes:** CORRESPONDING AUTHOR Dimitrios Pantazis, McGovern Institute for Brain Research, Massachusetts Institute of Technology Cambridge, Massachusetts, USA.

## Abstract

Visual gamma oscillations have been proposed to subserve perceptual binding, but their strong modulation by diverse stimulus features confounds interpretations of their precise functional role. Overcoming this challenge necessitates a comprehensive account of the relationship between gamma responses and stimulus features. Here we used multivariate pattern analyses on human MEG data to characterize the relationships between gamma responses and one basic stimulus feature, the orientation of contrast edges. Our findings confirmed we could decode orientation information from induced responses in two dominant frequency bands at 24-32 Hz and 50-58 Hz. Decoding was higher for cardinal than oblique orientations, with similar results also obtained for evoked MEG responses. In contrast to multivariate analyses, orientation information was mostly absent in univariate signals: evoked and induced responses in early visual cortex were similar in all orientations, with only exception an inverse oblique effect observed in induced responses, such that cardinal orientations produced weaker oscillatory signals than oblique orientations. Taken together, our results showed multivariate methods are well suited for the analysis of gamma oscillations, with multivariate patterns robustly encoding orientation information and predominantly discriminating cardinal from oblique stimuli.

## 2 Introduction

To yield a coherent perception of the visual world, the human brain must bind simple features into complex wholes. While substantial evidence underscores the importance of the rhythmic firing of neurons in the gamma band (30-80 Hz) for feature binding (Gray, Charles, 1989; Engel et al., 1991; Singer, 1999; Tallon-Baudry and Bertrand, 1999; Fries et al., 2007), the specific mechanisms subserving the temporal linking of visual information remain unclear (Shadlen and Movshon, 1999; Brunet et al., 2014; Ray and Maunsell, 2015).

A principal challenge in tackling this problem is understanding the modulation of visual gamma activity with respect to different stimulus features. A growing number of electrophysiological studies suggest both visual gamma power and frequency are strongly and systematically modulated by diverse stimulus features, including spatial frequency (Adjamian et al., 2004; Hadjipapas et al., 2007), luminance contrast (Adjamian et al., 2004; Hall et al., 2005), velocity (Swettenham et al., 2009), orientation (Koelewijn et al., 2011), size (Muthukumaraswamy and Singh, 2013), and eccentricity (van Pelt and Fries, 2013). The mounting list of features that can modulate gamma oscillations complicates interpretations on the role of visual gamma activity in feature binding.

Here, we propose the use of multivariate pattern analyses methods to obtain a fine characterization of the information encoded in gamma oscillations. The motivation is threefold. First, gamma power and frequency vary across the cortex, stressing the need to characterize gamma spatial patterns. For example, neuronal ensembles in different patches of cortex have been shown to oscillate at significantly different frequencies in response to stimuli with spatially varying contrast (Ray and Maunsell, 2010). Second, the strong dependence of gamma activity on diverse stimulus features implies complex relationships that cannot be captured by only two variables. Thus it is not possible to fully characterize the complexity of gamma signals using univariate approaches, which summarize activity across large patches of cortex into two values, the gamma power and frequency within the cortical patch (Adjamian et al., 2008; Koelewijn et al., 2011; Muthukumaraswamy and Singh, 2013). And third, multivariate pattern analyses methods have already been proven effective in resolving information encoded in MEG and EEG evoked responses, motivating their extension to induced responses. In particular, recent studies have revealed that information on simple visual features, such as orientation of contrast edges (Cichy et al., 2015; Ramkumar et al., 2013; Wardle et al., 2016), and complex visual patterns, such as visual object categories (Carlson et al., 2013; Cichy et al., 2014; Isik et al., 2014; Clarke et al., 2014; Kaneshiro et al., 2015), are encoded in MEG and EEG evoked responses.

We used multivariate analyses methods to systematically evaluate the role of gamma oscillations in encoding a simple stimulus feature, the orientation of grating stimuli (Koelewijn et al., 2011). We recorded human MEG data while participants viewed six circular grating stimuli with different orientations (0-150° in 30° steps), and used multivariate pattern classification on MEG sensor measurements to decode stimulus orientation from induced responses in a broad range of frequencies (10-80 Hz). The selection of stimuli enabled us to compare brain responses to orientations differing as little as 30°. We also evaluated whether induced responses discriminate cardinal (0°, 90°) from oblique orientations (30°, 60°, 120°, 150°), as predicted by behavioral, electrophysiological and neuroimaging studies of the oblique effect (Pettigrew et al., 1968; Appelle, 1972; Koelewijn et al., 2011). Finally, we related findings on induced responses with corresponding multivariate analyses of MEG evoked responses.

## 3 Methods

### 3.1 Participants

Fourteen right handed healthy subjects (8 females; age mean ± s.d. = 27.2 ± 5.7 years) participated in the experiment. All subjects gave a written informed consent and received payment for their participation. The study was approved by the local ethics committee (Institutional Review Board of the Massachusetts Institute of Technology) and conducted according to the principles of the Declaration of Helsinki.

### 3.2 Stimulus set and experimental design

The stimulus set comprised 6 stationary square-wave Cartesian gratings with orientations 0° to 150° with respect to vertical, in steps of 30° (Fig. 1A). This enabled the evaluation of brain responses to orientations differing by as little as 30°, thus establishing a measure of sensitivity in disambiguating orientations. It also allowed the comparison of specific orientation combinations, namely cardinal (0°, 90°) versus oblique (30°, 60°, 120°, 150°).

**Fig. 1.**
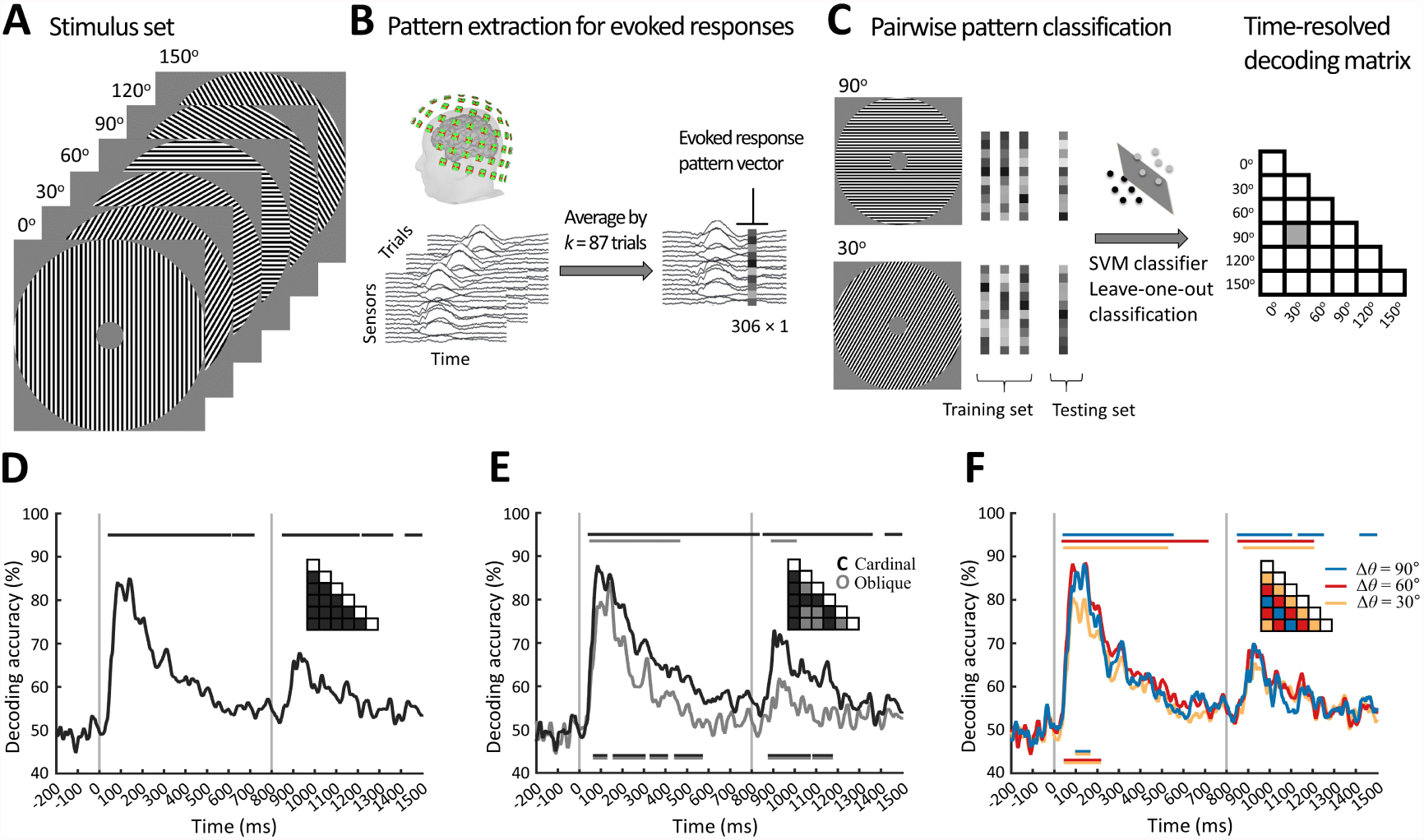
Orientation decoding from MEG evoked responses. **A)** The stimulus set comprised 6 Cartesian square-wave gratings with orientations 0° to 150° with respect to vertical, in steps of 30°. **B)** Construction of MEG evoked response pattern vectors. Condition-specific MEG trials were first averaged by *k* = 87 trials to increase SNR, and then sensor measurements for a given time point t were concatenated in 306 dimensional time-resolved pattern vectors. **C)** Multivariate pattern analysis. For each time point *t*, a support vector machine (SVM) was trained to discriminate pairs of stimuli (here exemplified for 90° and 30°), and the resulting pairwise decoding accuracies populated the elements of a 6 × 6 decoding matrix. **D)** Decoding accuracy averaged across all pairs of stimuli. **E)** Decoding accuracy for pairs of stimuli when both have oblique orientations (gray), or at least one of them has a cardinal orientation (black). **F)** Decoding accuracy for pairs of stimuli with orientation angle disparities Δ*θ* of 30°, 60°, and 90°. For onset and peak latencies see Table 1A. Gray vertical lines indicate stimulus onset and offset. Inset matrices indicate which matrix segments were averaged, color-coded as in decoding curves. Lines above plots indicate significant time points, color-coded as in decoding curves. Double-colored lines below plots indicate significant differences between decoding time series of corresponding colors (*N* = 14; sign permutation test; one- or two-sided for decoding time series and their differences, respectively; *p* < 0.05 cluster defining threshold; *p* < 0.05 cluster threshold).

The gratings had black/white maximum contrast with a frequency of 3 cycles per degree, which is known to elicit strong evoked responses and induced narrow band gamma oscillations in the early visual cortex (Adjamian et al., 2004; Koelewijn et al., 2011). The stimuli were constrained in an annulus with an outer radius of 6.5° and an inner radius of 1°. The inner radius served to prevent interaction effects between the orientation contrast edges and a fixation cross presented at the center of the stimulus during the experiment.

The grating stimuli were presented against a gray background, while the central fixation cross remained always on screen (Suppl. Fig. 1). Images appeared in random order for 800 ms with an ISI of 1000 ms. The gratings had random phase in each trial to ensure any identified orientation representations are not confounded by local luminance differences due to a particular choice of phase (Ramkumar et al., 2013; Cichy et al., 2015). Every 3-5 trials the sequence was interrupted to present a target image (paper clip) for 800 ms followed by a longer ISI of 1500 ms. The target trials served to maintain attention and avoid contamination of experimental conditions with eye blink artifacts, and were excluded from further analysis.

The experiment consisted of 22 blocks of 96 trials each (excluding the interspersed targets), for a total of 352 trials per stimulus. Participants were instructed to fixate on a centrally presented black fixation cross, and press a button and blink their eyes in response to the target image. The stimuli were presented using Psychtoolbox (www.psychtoolbox.org) (Brainard, 1997).

### 3.3 MEG acquisition and preprocessing

MEG data was recorded using an Elekta Triux system (306-channel probe unit with 204 planar gradiometer sensors and 102 magnetometer sensors) at a sampling rate of 1000 Hz, filtered between 0.03 and 330 Hz. The location of the head was measured continuously during each recording session by activating a set of 5 head position indicator coils placed over the head. Prior to the MEG recording, a 3D digitizer (Fastrak, Polhemus, Colchester, Vermont, USA) was used to register the locations of 3 anatomical landmarks (right and left preauricular points and the nasion) with respect to the 5 head position indicator coils. Raw data was pre-processed with the Maxfilter software (Elekta, Stockholm) to compensate for head movements and perform noise reduction with spatiotemporal filters (Taulu et al., 2004; Taulu and Simola, 2006). We used default parameters (harmonic expansion origin in head frame = [0 0 40] mm; expansion limit for internal multipole base = 8; expansion limit for external multipole base = 3; bad channels automatically excluded from harmonic expansions = 7 s.d. above average; temporal correlation limit = .98; buffer length = 10 s). Intuitively, Maxfilter first applied a spatial filter that separated the signal data from spatial patterns emanating from distant noise sources outside the sensor helmet. It then applied a temporal filter that discarded components of the signal data with time series strongly correlated with the ones from the noise data.

The filtered data were subsequently analyzed with Brainstorm (Tadel et al., 2011). We extracted peri-stimulus data from −200 ms to +1500 ms with respect to stimulus onset. Every trial was baseline-corrected to remove the mean (−300 to 0 ms) from each channel, and the time series were smoothed with a 30 Hz low-pass filter for evoked analyses, or 200 Hz low-pass filter for induced response analyses. For multivariate pattern analysis, in addition to removing the baseline mean, the data of each sensor were divided by the standard deviation of the pre-stimulus baseline signal of that sensor.

### 3.4 Multivariate pattern analysis of evoked responses

We first evaluated whether stimulus orientation can be decoded from evoked responses (Fig. 1). Multivariate pattern analysis was based on linear support vector machines (SVM) using the libsvm software implementation (Chang and Lin, 2011) with a fixed regularization parameter *C* = 1. To reduce computational load and improve the signal-to-noise ratio, we sub-averaged the *M* = 352 trials per condition in groups of *k* = 87 trials with random assignment, obtaining 4 averaged trials per condition (Fig. 1B).

SVM analysis was performed separately for each subject in a time-resolved manner. In particular, for each time point *t* (from -200 ms to 1500 ms in 1 ms steps), we extracted evoked pattern vectors by concatenating the 306 MEG sensor measurements into 306-dimensional pattern vectors, resulting in 4 pattern vectors for each stimulus (condition). For each pairwise combination of conditions separately, we measured the performance of the classifier to discriminate between conditions using leave-one-out cross-validation: 3 vectors were randomly assigned to the training set, and the left-out vector to the testing set to evaluate the classifier decoding accuracy. The pairwise classification was repeated 100 times with random assignments of the *M* trials in groups of *k* trials, and the resulting decoding accuracies were averaged over repetitions. This yielded a 6 × 6 decoding matrix for each time point *t*, indexed in rows and columns by the classified stimuli. This decoding matrix is symmetric and has an undefined diagonal (no classification within condition).

Based on the decoding results, we further determined whether MEG activation patterns discriminate cardinal from oblique orientations, and whether higher orientation disparities are associated with greater decoding accuracies. We thus partitioned the decoding matrix into segments for pairs of stimuli when both have oblique orientations (segment O), or at least one of them has a cardinal orientation (segment C). We also separately partitioned the decoding matrix into 3 segments, corresponding to pairs of stimuli with orientation angle disparities Δ*θ* of 30°, 60°, and 90°. Averaging decoding accuracies within each of the segments enabled the comparison of the corresponding activation patterns.

### 3.5 Multivariate pattern analysis of induced responses

For induced response analysis, the evoked response of each condition (trial average) was subtracted from the individual *M* = 352 trials of that condition (Fig. 2A). For each trial and sensor, the MEG time series were then transformed to time-frequency power maps in the range 10 – 80 Hz by convolving them with complex Morlet wavelets described by the equation:

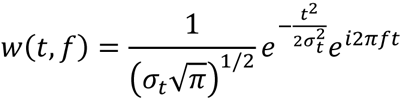

**Fig. 2.**
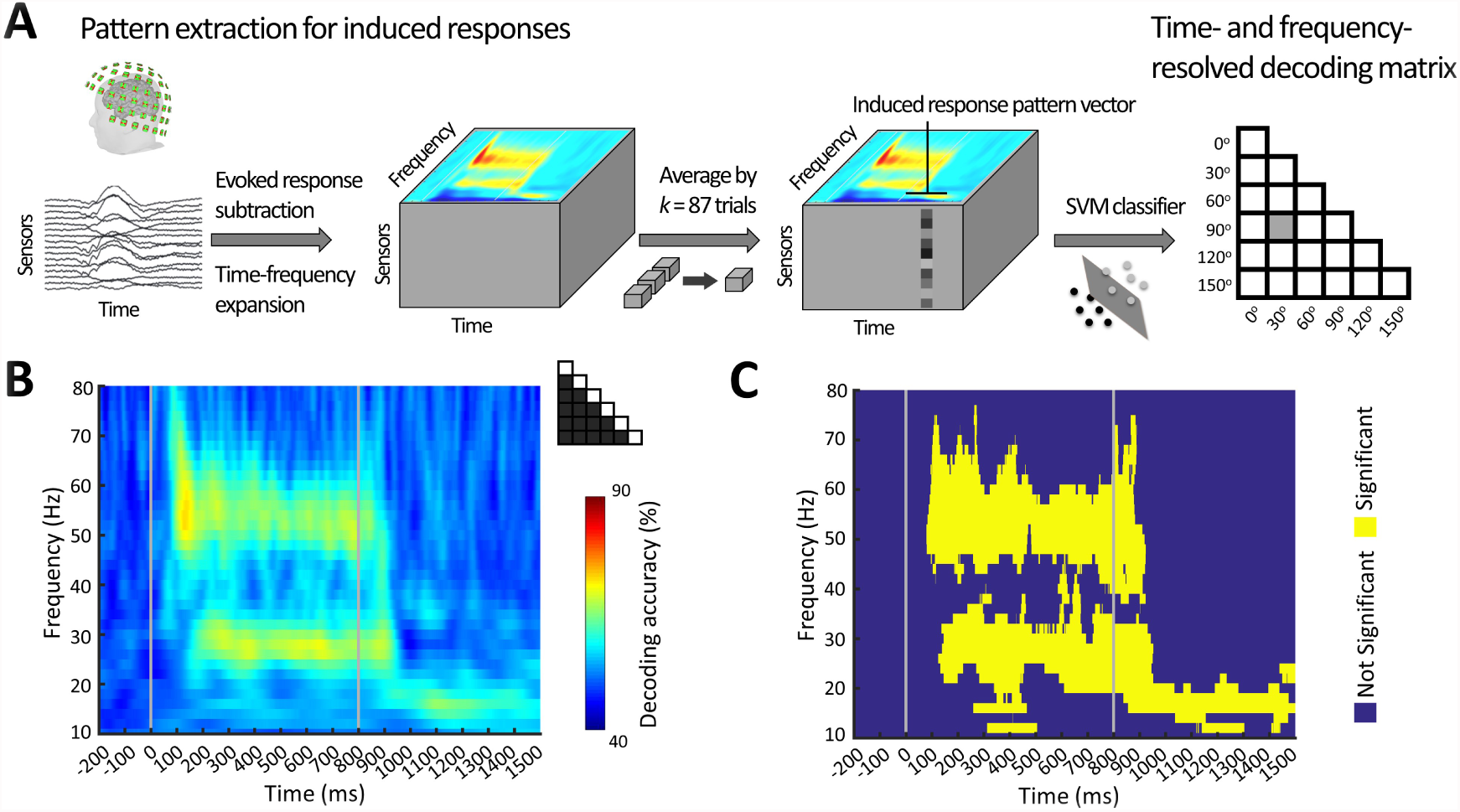
Orientation decoding from MEG induced responses. **A)** Construction of induced response MEG pattern vectors. Following removal of the evoked response, condition-specific single trial MEG data were expanded in time-frequency components using complex Morlet wavelets. The resulting data were averaged by *k* = 87 trials, and the sensor measurements were concatenated in 306 dimensional time- and frequency-resolved induced pattern vectors. For each time and frequency points, the induced response pattern vectors were subjected to multivariate classification analysis analogous to Fig 1C, resulting in time- and frequency-resolved decoding matrices. **B)** Time-frequency map of decoding accuracies. Results were averaged across all pairs of stimuli as indicated by the inset matrix. **C)** Statistical significance map (*N* = 14; one-sided sign-permutation test; *p* < 0.005 cluster defining threshold; *p* < 0.05 cluster threshold). Gray vertical lines indicate stimulus onset and offset.

The complex Morlet wavelets have Gaussian shape in both the time and frequency domain, with a temporal resolution 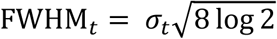 and a spectral resolution 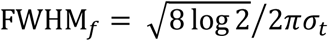, where *σ_t_* is the standard deviation of the Gaussian in the time domain. The wavelets were selected to have a constant ratio FWHM_*t*_/*f* = 3, which for example corresponds to temporal resolution FWHM_*t*_ = 3 s at central frequency *f* = 1 Hz.

The resulting *M* = 352 time-frequency maps per condition were averaged in groups of *k* = 87 with random assignment to reduce computational load, improve the signal-to-noise ratio, and allow direct comparison with the similarly analyzed evoked responses. This yielded 4 averaged time-frequency maps per condition (Fig. 2A). These maps were finally normalized as a percentage change from the average baseline power for each frequency band, to represent event related synchronization/desynchronization (ERS/ERD).

SVM analysis was performed in a time- and frequency resolved manner. In particular, for each time point *t* (from -200 ms to 1500 ms in 1 ms steps), and frequency value *f* (from 10 Hz to 80 Hz in 2 Hz steps) we extracted induced pattern vectors by concatenating the sensor time-frequency values into 306-dimensional pattern vectors. This procedure yielded 4 pattern vectors for each condition. Pairwise SVM classification proceeded similarly to the evoke response analyses, by performing 100 iterations of leave-one-out cross-validation (thus using 3 pattern vectors to construct the SVM hyperplane and 1 pattern vector to test the prediction of the classifier), yielding a 6 × 6 decoding matrix for each time point t and frequency point *f* (Fig. 2A).

To further characterize the decoding performance in 2 frequency bands, 50-58 Hz (high gamma) and 24-32 Hz (low gamma) (Fig. 3), we designed time-resolved higher dimensional pattern vectors by concatenating the power maps values from 306 sensors and 5 frequency points (50-58 Hz or 24-32 Hz, in 2 Hz steps), resulting in 1530 × 1 dimensional pattern vectors. Other than using these higher dimensional induced response pattern vectors, multivariate pattern analysis proceeded identically to the previously described procedures.

**Fig 3.**
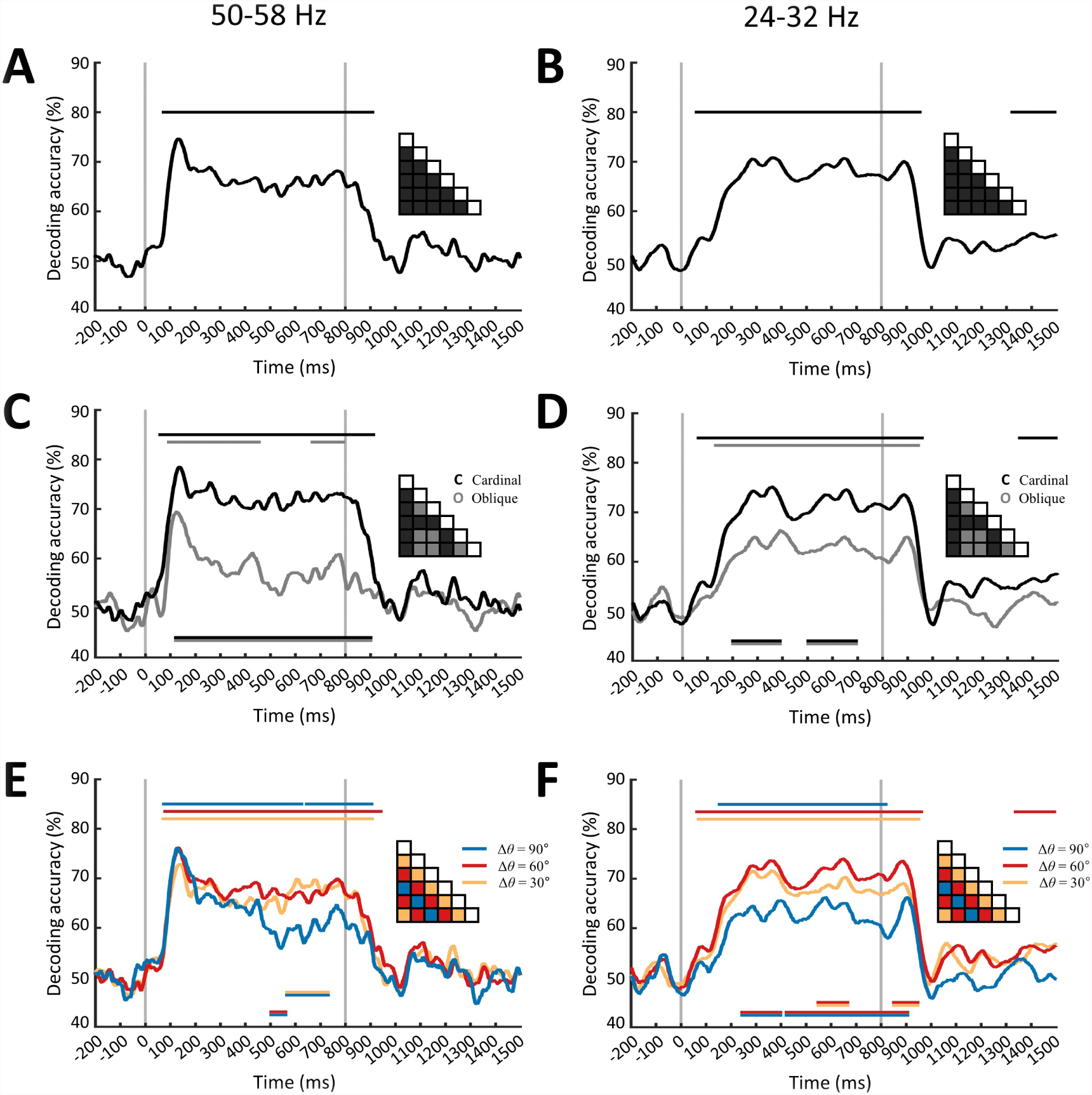
Orientation decoding from MEG induced responses at 50-58Hz and 24-32Hz. **AB**) Decoding accuracy averaged across all pairs of stimuli. **CD**) Decoding accuracy for pairs of stimuli when both have oblique orientations (gray), or at least one of them has a cardinal orientation (black). **F**) Decoding accuracy for pairs of stimuli with orientation angle disparities Δ*θ* of 30°, 60°, and 90°. Gray vertical lines and lines above and below plots are same as in Fig 1 (*N* = 14; sign permutation test; one- or two-sided for decoding time series and their differences, respectively; *p* < 0.05 cluster defining threshold; *p* < 0.05 cluster threshold)

To evaluate cardinal vs. oblique effects, and the discrimination across several angle disparities Δ*θ*, the 6 × 6 induced response decoding matrices were partitioned into appropriate segments (Fig. 3C-F), reflecting the corresponding evoked response analysis.

### 3.6 Temporal generalization of multivariate pattern analysis

The distinct temporal dynamics of the transient evoked time series versus the sustained induced gamma responses of the early visual cortex posit the question whether orientation information is maintained in different ways between the two responses. To evaluate the persistence of orientation information over time, we generalized the decoding procedure across time by training the SVM classifier at a given time point *t*, as before, but testing across all other time points (Cichy et al., 2014; King and Dehaene, 2014; Isik et al., 2014). Intuitively, if representations are stable over time, the classifier should successfully discriminate signals not only at the trained time *t*, but also over extended periods of time that share the same neural representation of orientation information. This temporal generalization analysis was repeated for every pair of stimuli, and the results were averaged across conditions and subjects, yielding 2-dimensional temporal generalization matrices with the x-axis denoting training time and the y-axis testing time (Fig. 4).

**Fig. 4.**
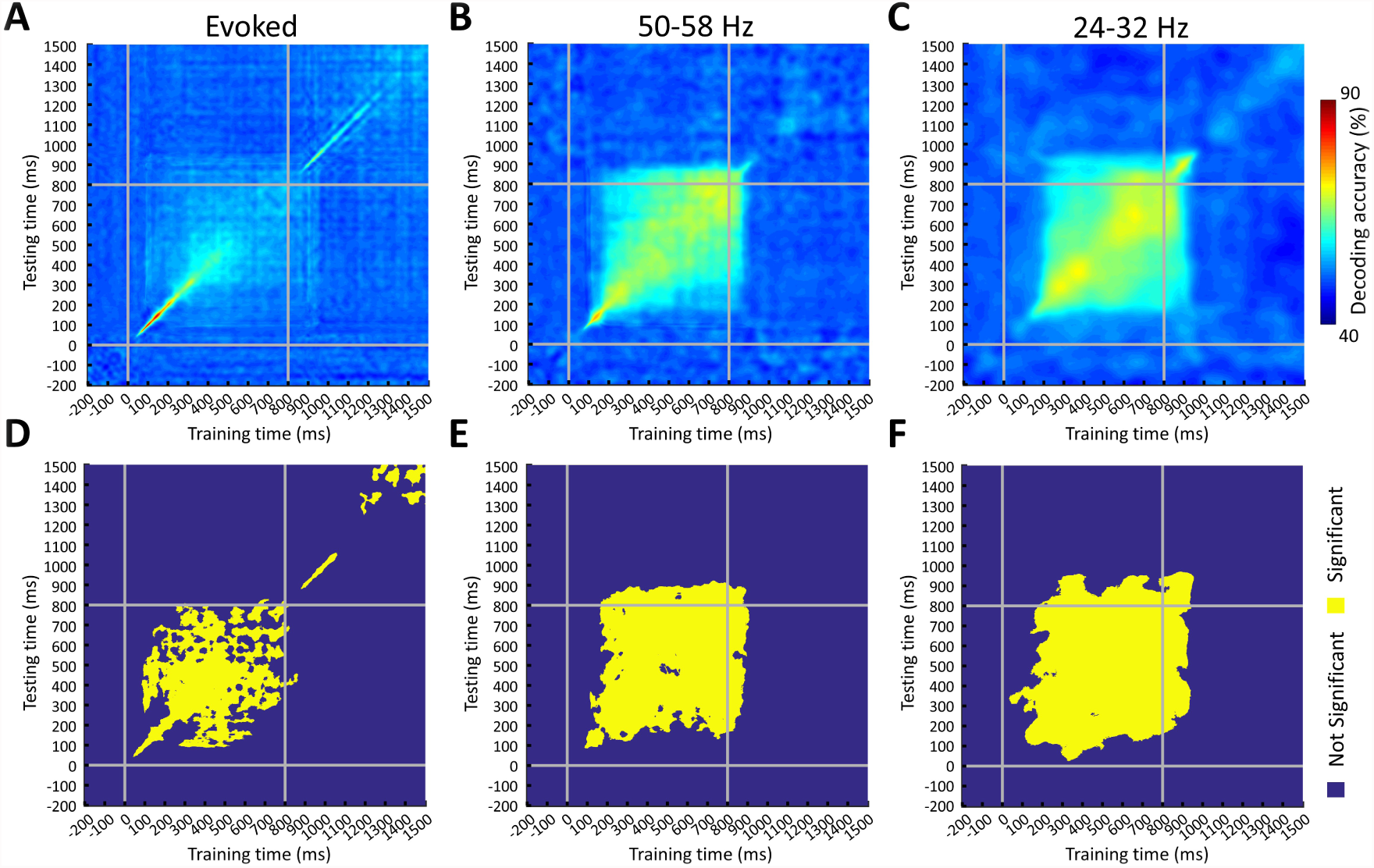
Temporal generalization of orientation decoding from evoked and induced 50-58 Hz and 24-32 Hz MEG signals. A support vector machine classifier was trained with MEG brain responses at any given time point t (x-axes) and tested against all other time points (y-axes). **A)** Evoked response temporal generalization map. Decoding accuracy is averaged across all pairs of stimuli and subjects. **BC)** Same as A but for induced response at 50-58 Hz and 24-32 Hz. **DEF)** Statistical significance maps for corresponding temporal generalization matrices (*N* = 14; one-sided sign-permutation test; *p* < 0.005 cluster defining threshold; *p* < 0.05 cluster threshold). Gray vertical and horizontal lines indicate stimulus onset and offset

### 3.7 Comparison of evoked and induced representations to models

The MEG decoding matrices can be interpreted as a dissimilarity measure: stimulus pairs with similar neural patterns are harder to discriminate and thus have lower decoding accuracies. Termed representational dissimilarity matrices (RDMs), these 6 × 6 decoding matrices capture the relations across the neural patterns elicited by the 6 stimulus orientations.

To evaluate the orientation information encoded in the evoked and induced RDMs, we devised two models reflecting hypothesized representational formats (Fig. 5A). The cardinal model was a 6 × 6 matrix with representational distance 2 for pairs of stimuli with at least one of them cardinal, and 1 for pairs of stimuli with both oblique. This model evaluated a categorical relationship between cardinal and oblique stimuli. The angle disparity model was a 6 × 6 matrix with representational distance equal to the angle disparity Δ*θ* between the corresponding pairs of stimuli. This model thus assumed an ordinal relationship between angle disparities and neuronal representations, with higher angle disparities associated with increasingly different neuronal representations.

**Fig. 5.**
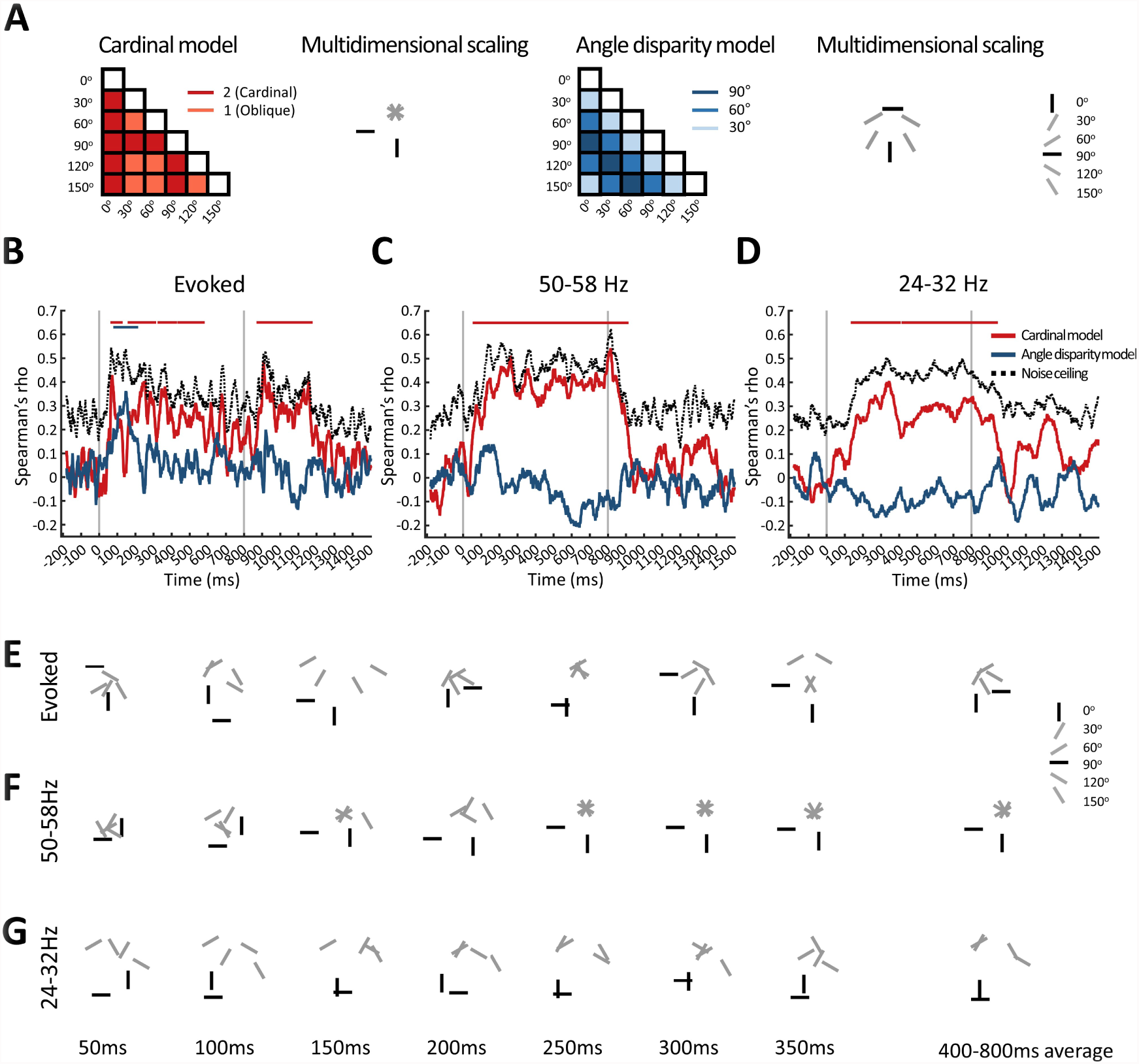
Comparison of MEG evoked and induced responses to hypothesized models of orientation encoding. **A)** Cardinal and angle disparity models shown with their MDS plots. **B)** Representational similarity of the MEG evoked responses with the two models. For every time point, the MEG decoding matrix was rank correlated with each of the models. Black dashed lines denote the noise ceiling. Gray vertical lines and lines above plots same as in Fig 1 (*N* = 14; one-sided sign permutation test; *p* < 0.05 cluster defining threshold; *p* < 0.05 cluster threshold). **CD)** Same as A for induced 50-58 Hz and 24-32 Hz responses. **EFG)** MDS plots for evoked and induced responses sampled every 50 ms.

We then compared (using Spearman’s rho to capture ordinal relationships) the two model RDMs with the time-resolved MEG RDMs of the evoked response and the induced response in the high and low gamma bands (Fig. 5B). This resulted in time courses of representational similarity between the models and MEG data.

To further characterize the effectiveness of these models in explaining MEG orientation information, we computed the corresponding noise ceiling in each time point, defined as the highest possible correlation attained by the (unknown) true model, given the inherent noise in the MEG data (Nili et al., 2014). Intuitively, the model-MEG correlation is maximized when the model RDM lies exactly at the center of the cloud of single-subject MEG RDM estimates (in a rank-transformed space to account for Spearman’s correlation). Thus operationally, the noise ceiling was computed by rank-transforming the single-subject MEG RDMs for each time point *t*, and defining the subject-averaged MEG RDM as the ideal true model for that given point *t*.

### 3.8 Multidimensional scaling visualization

To offer an intuitive visualization of the complex representational patterns contained in the MEG and model RDMs, we used multidimensional scaling (MDS) (Fig. 5). MDS is an unsupervised method to visualize the level of similarity between individual conditions contained in a distance matrix. We plotted the data in a two-dimensional space, whereby similar orientations are grouped together and dissimilar orientations far apart. To ease visualization, the stimuli were represented by bars (black for cardinal; gray for oblique) with orientation identical to that of corresponding grating stimulus.

### 3.9 Orientation selective neuronal activation in early visual cortex

While multivariate pattern analyses and representational similarity can capture orientation information encoded in neuronal signals, they constitute distance measures between brain signals and thus abstract from the underlying brain activity. To directly characterize neuronal activation in early visual cortex, we computed source activation maps on subject-specific cortical surfaces derived from Freesurfer automatic segmentation (Fischl et al., 2004). The forward model was calculated using an overlapping spheres model (Huang et al., 1999). MEG signals were then mapped on the cortex using a dynamic statistical parametric mapping approach (dSPM) (Dale et al., 2000) and time series were derived from pericalcarine cortex (Desikan et al., 2006).

Evoked and induced neuronal activations in early visual cortex were computed similarly to sensor space, by averaging condition-specific MEG time series and using complex Morlet wavelets to estimate induced signals.

### 3.10 Statistical testing

We used non-parametric statistical inference that does not make assumptions on the distributions of the data (Maris and Oostenveld, 2007; Pantazis et al., 2005). Permutation and bootstrap tests were performed with sample size *N* = 14, equal to the number of subjects for random effects inference.

Statistical assessment of classification time series and matrices, MEG-model correlations, and brain activation matrices relied on sign permutation tests. The null hypothesis was equal to 50% chance level for decoding results, and 0 for correlation values, brain neuronal activations or power percentage change from baseline. In all cases, under the null hypothesis we could permute the condition labels of the MEG data, which was equivalent to a sign permutation test that randomly multiplied subject responses by +1 or -1. Repeating the permutation procedure 1000 times enabled us to convert the original statistical maps (e.g. decoding time series, MEG-model correlation time series) to 1-dimensional or 2-dimensional p-value statistical maps. We then controlled the familywise error across time points using cluster size inference. The p-value statistical maps were thresholded at *p* < 0.05 for 1-dimensional data, or *p* < 0.005 for 2-dimensional data, to define suprathreshold clusters. These suprathreshold clusters were reported as significant if their size (defined as the number of connected elements) exceeded a *p* < 0.05 threshold, defined by the empirical distribution of similarly constructed suprathreshold clusters of the permutation sample statistical maps.

Statistical assessment of brain activation time series also relied on sign permutation tests to convert statistics into p-values, however multiple comparison corrections over time were controlled with a false discovery rate (FDR) procedure (*p* < 0.05) because it was more suitable than the cluster size inference, given the highly transient temporal dynamics.

Statistical tests were one-sided for decoding values, and two-sided for decoding value differences or brain activation values and differences.

Statistical assessment of the peak latency of the time series relied on bootstrap tests to estimate confidence intervals. The subject-specific time series were bootstrapped 1000 times and the empirical distribution of the peak latency of the subject-averaged time series was used to define 95% confidence intervals. For peak-to-peak latency differences, we obtained 1000 bootstrapped samples of each of the two peaks, and their statistical difference was evaluated with a two-sample two-sided signed-rank test.

## 4 Results

### 4.1 Orientation information is decodable from MEG evoked responses

We first determined the time series with which evoked responses detected by MEG discriminate orientation information. For every time point, we averaged all elements of the MEG decoding matrix, yielding a grand total decoding time series averaged across all condition pairs (Fig. 1D). MEG evoked responses robustly resolved orientation information, reproducing the results in (Cichy et al., 2015; Ramkumar et al., 2013). The decoding time series reached significance at 38 ms (95% confidence interval 37 – 39 ms), peaked at 141 ms (83 – 145 ms), and then progressively declined over time.

Next we investigated whether MEG evoked responses differentially encode cardinal (0°, 90°) from oblique (30°, 60°, 120°, 150°) orientations. Such division is behaviorally relevant, given the perceptual differences in orientation discrimination between cardinal (i.e. vertical and horizontal) and oblique contours (Appelle, 1972). While this oblique effect may be a consequence of living in natural and manufactured environments with straight horizontal and vertical lines dominating our surroundings (Torralba and Oliva, 2003), the neural underpinnings remain controversial. In part, this is because the brain responses may be more complex than a simple bias (also see horizontal bias; Essock et al., 2003), and controversial findings are possibly driven by disparate methodological approaches (Brunet et al., 2014; Dora Hermes et al., 2015; Maloney and Clifford, 2015).

To investigate differences in the encoding of cardinal vs. oblique orientations anisotropies, we partitioned the MEG decoding matrix into 2 segments; a cardinal segment containing elements with pairs of stimuli when at least one of them is cardinal (segment C; black); and an oblique segment when both stimuli are oblique (segment O; gray). After averaging decoding accuracies within the corresponding segments, we observed higher decoding accuracies for cardinal than oblique stimuli pairs (Fig. 1E). The effect was consistent throughout the evoked response, with the preference for cardinal stimuli extending even during the offset response.

To evaluate the effect of angle disparities on the decoding time series, we further partitioned the decoding matrix into 3 segments corresponding to pairs of stimuli with orientation differences Δ*θ* of 30°, 60°, and 90°. While differences even as low as 30° were decodable, in contrast to the consistent preference to cardinal stimuli, angle disparity effects were less evident in the MEG evoked responses (Fig. 1F). The decoding accuracies for the 30° case were weaker than the 60° and 90° cases only during the first couple hundred milliseconds after the stimulus onset, with responses equivalent thereafter. Onset and peak latencies for the grand total, cardinal vs. oblique, and the angle disparities of 30, 60, and 90 are available in Table 1A.

**Table 1.** Onset and peak latency of orientation decoding time series for A) evoked response, B) induced response 50-58 Hz, and C) induced response 24-32 Hz. Numbers in brackets indicate 95% confidence intervals.

|  | Onset latency (ms) | Peak latency (ms) |
| --- | --- | --- |
| **A) Evoked** |  |  |
| Grand total | 38 (37 – 39) | 141 (83 – 145) |
| Cardinal | 40 (39 – 40) | 86 (81 – 143) |
| Oblique | 46 (45 – 47) | 141 (88 – 145) |
| Angle disparity 90° | 39 (38 – 39) | 139 (98 – 149) |
| Angle disparity 60° | 35 (34 – 36) | 145 (82 – 148) |
| Angle disparity 30° | 42 (41 – 42) | 86 (83 – 142) |
| **B) 50-58 Hz** |  |  |
| Grand total | 68 (64 – 70) | 134 (122 – 154) |
| Cardinal | 51 (48 – 52) | 138 (130 – 750) |
| Oblique | 86 (86 – 87) | 126 (113 – 154) |
| Angle disparity 90° | 68 (65 – 73) | 128 (123 – 332) |
| Angle disparity 60° | 72 (71 – 73) | 134 (118 – 443) |
| Angle disparity 30° | 62 (61 – 68) | 138 (124 – 775) |
| **C) 24-32 Hz** |  |  |
| Grand total | 58 (54 – 60) | 368 (278 – 903) |
| Cardinal | 51 (49 – 59) | 360 (270 – 903) |
| Oblique | 127 (122 – 134) | 391 (208 – 907) |
| Angle disparity 90° | 143 (141 – 150) | 579 (286 – 911) |
| Angle disparity 60° | 53 (51 – 58) | 649 (282 – 904) |
| Angle disparity 30° | 62 (61 – 66) | 361 (266 – 901) |

Overall, we found that MEG evoked responses robustly encode orientation information during the first few hundred milliseconds, with consistently higher discrimination accuracy for cardinal than oblique orientations. There was limited evidence for angle disparity effects; all angle disparities were equally decodable, with only exception a weakened discrimination of small disparities (30°) the first couple hundred milliseconds after the stimulus onset.

### 4.2 Orientation information is decodable from induced responses

Grating stimuli are known to elicit strong induced gamma activity in the visual cortex, with amplitude and peak frequency dependent on stimulus features, such as spatial frequency, contrast, and velocity (Adjamian et al., 2004; Friedman-Hill et al., 2000). Here we investigated whether induced MEG responses, particular in the gamma range, also encode grating orientation information. We computed MEG induced responses in the range 10 – 80 Hz, and used multivariate pattern analysis to obtain time- and frequency-resolved MEG decoding matrices for the 6 square-wave Cartesian gratings (Fig. 2A).

Averaging all elements of the decoding matrices yielded time-frequency maps of grand total decoding accuracy (Fig. 2BC). MEG induced responses demonstrated strong orientation selectivity in two frequency bands: 50-58 Hz and 24-32 Hz. The 50-58 Hz frequency band is the narrow-band induced gamma activity consistently reported in the literature in response to visual stimulus gratings (Hadjipapas et al., 2015; Koelewijn et al., 2011; Muthukumaraswamy and Singh, 2013; Perry et al., 2013). We found the 24-32 Hz frequency band to also carry orientation information with decoding performance comparable to the 50-58 Hz frequency band. In both cases, decoding was sustained for the entire 800 ms duration of stimulus presentation.

We further investigated higher frequencies up to 190 Hz with the same decoding procedure. Other than the two frequency bands, we found no orientation selectivity for higher frequencies (Suppl. Fig. 2). Furthermore, the two frequency bands were related to each other, with the 24-32 Hz band being a subharmonic of the 50-58 Hz band (Suppl. Fig. 3). This was determined by estimating the peak frequency for the two bands separately per subject and computing the ratio of high/low frequency peak. This ratio had an average value of 1.96 which was not statistically different from 2 (p = 0.31; two-sided signed-rank test against 2).

In sum, multivariate analyses captured orientation information in two frequency bands, 50-58 Hz and the subharmonic 24-32 Hz, providing the basis for an in-depth investigation of orientation encoding in these frequency bands.

### 4.3 Orientation selectivity of the MEG 50-58 Hz and 28-32 Hz induced responses

We further investigated the characteristics of the orientation information encoded in induced responses by focusing on the two identified and pertinent frequency bands: 24-32 Hz and 50-58 Hz. To this goal we refined the multivariate analysis procedure by constructing higher dimensional (1530 ×1) induced response pattern vectors, with the 5 · 306 = 1530 elements comprising time-resolved power measurements at 306 MEG sensors for 5 frequency values (50-58 Hz or 24-32 Hz, in 2-Hz steps).

Pairwise pattern classification resulted in 6 × 6 time-resolved decoding matrices for each frequency band, and averaging across all elements of the matrices yielded grand total decoding time series for the 50-58 Hz and 24-32 Hz bands (Fig. 3AB). Following an early transient response, both frequency bands reached a plateau of sustained 60-65% decoding that lasted until the offset of the stimuli. However, the shape of the transient response differed between the two bands, with the 50-58 Hz band reaching an early peak of 134 ms (122 – 154 ms) compared to the progressive increase and late peak of 368 ms (278 – 903 ms) for the 24-32 Hz band (see Table 1BC). The difference between the two peaks was statistically significant (Δ = 234 ms; *p* < 0.0001 two-sided signed-rank test).

We also determined the time series with which the induced responses resolved cardinal, oblique, and 30°, 60°, and 90° angle disparities (Fig 3C-F). Similarly to the analysis of evoked responses, we partitioned the MEG decoding matrices into segments and averaged the corresponding decoding accuracies. Both 50-58 Hz and 24-32Hz bands had higher decoding accuracies for cardinal than oblique stimuli. Angle disparity effects were less consistent, though the angle disparity Δ*θ* = 90° resulted in consistently weaker decoding accuracies than the 60° and 30° cases.

In sum, we found both frequency bands had sustained orientation information with robust preference to cardinal orientations. The 50-58 Hz response peaked early at 134 ms, whereas the 24-32 Hz response had a significantly delayed peak at 368 ms.

### 4.4 Orientation information is sustained over time in both evoked and induced MEG signals

The stable decoding time series of the induced responses, sustained over the duration of stimulus presentation, predict persistent neuronal orientation selectivity over time. We tested this prediction by performing a temporal generalization of the multivariate pattern analysis. In particular, we trained an SVM classifier to decode oriented stimuli at a given time point *t*, and then tested classification performance across all other time points (Cichy et al., 2014; Isik et al., 2014; King and Dehaene, 2014). This resulted in 2-dimensional decoding matrices with the x-axis indexed by training time, and the y-axis by testing time.

The results of the temporal generalization analysis confirmed the prediction: induced responses carry the same orientation information throughout the 800 ms stimulus presentation (Fig. 4BCEF). The significance maps formed square shapes for both the 50-58 Hz and 24-32 Hz bands, a signature pattern of sustained neuronal activity (King and Dehaene, 2014). By contrast, the evoked response maps show an early narrow diagonal that subsequently broadens considerably, consistent with an early transient and then sustained pattern of activity (Fig. 4AD).

Taken together, the evoked and induced responses had distinct temporal dynamics. The temporal generalization of the evoked response, with a rapid evolution of neuronal activity early and a greater generalization relative late after stimulus onset, matched in shape corresponding results from other studies despite using diverse types of stimuli (Carlson et al., 2013; Cichy et al., 2014; Isik et al., 2014). In contrast, induced responses were predominantly sustained and generalized equally well throughout the neuronal activity.

### 4.5 Evoked and induced representations are largely explained by cardinal orientation selectivity

To comprehensively characterize the representational structure of the evoked and induced responses, we tested two alternative hypotheses motivated by the above decoding results: 1) we hypothesized the orientations follow a categorical representational structure solely explained by cardinal versus oblique orientations, and 2) we hypothesized there is an ordinal relationship between angle disparities and neuronal pattern differences, such that higher angle disparities were associated with more dissimilar neuronal representations. We devised two models to evaluate the hypothesized representations (Fig. 5A). A cardinal model had elements 2 for pairs of stimuli with at least one cardinal, and 1 otherwise. An angle disparity model had elements equal to the angle disparity Δ*θ* between the corresponding pairs of stimuli. MDS plots of these models, offering an intuitive visualization of the imposed similarity between the individual stimuli, are shown next to the respective models. In these plots, stimuli are depicted by bars (black for cardinal; gray for oblique) with orientation matching that of corresponding stimuli.

The two models were compared (Spearman’s rho) against the time-resolved MEG decoding matrices of the evoked and induced responses (Fig 5BCD). In all cases, the cardinal model predominantly explained the MEG data. Importantly, its performance was close to noise ceiling, i.e. the highest possible correlation attained by the (unknown) true model, given the inherent noise in the MEG data (Nili et al., 2014). Conversely, the angle disparity model proved a poor candidate model, with weak overall correlations. There was one case where the angle disparity model reached significance, even exceeding the cardinal model, during the range 74 - 216 ms in the evoked response.

We further created MDS plots of the MEG responses for intuitive visualization, sampled every 50 ms (Fig. 5EFG). As representations evolved over time, the cardinal selectivity became apparent in all cases, with the black bars clustering away from the remaining gray. A marked example is the 50-58Hz induced response, with MDS results practically matching the cardinal model.

Overall, the cardinal model offered an accurate characterization of the induced response orientation representations, with performance near noise ceiling. It was also an effective descriptor of orientation representations in evoked responses, though early MEG evoked signals were better explained by an angle disparity model.

### 4.6 Orientation selectivity of neuronal activation in early visual cortex

While multivariate pattern analysis resolved orientation information, decoding accuracies abstracted away from the underlying neuronal activation, and thus did not directly inform us how neural activities differed. Similarly, representational similarity constitutes a second-order description of the MEG data, documenting relations between stimulus-specific neural activity patterns rather than the patterns themselves. To directly characterize neuronal activation in cortex, we mapped MEG sensor maps to cortical sources using a dSPM approach (Dale et al., 2000). Cortical maps of MEG induced responses at 50-58Hz and 24-32Hz are shown in Fig. 6. Visual gamma oscillations for both frequency bands localized on early visual cortex (EVC), consistent with prior findings (Adjamian et al., 2004; Hoogenboom et al., 2006; Koelewijn et al., 2011).

**Fig. 6.**
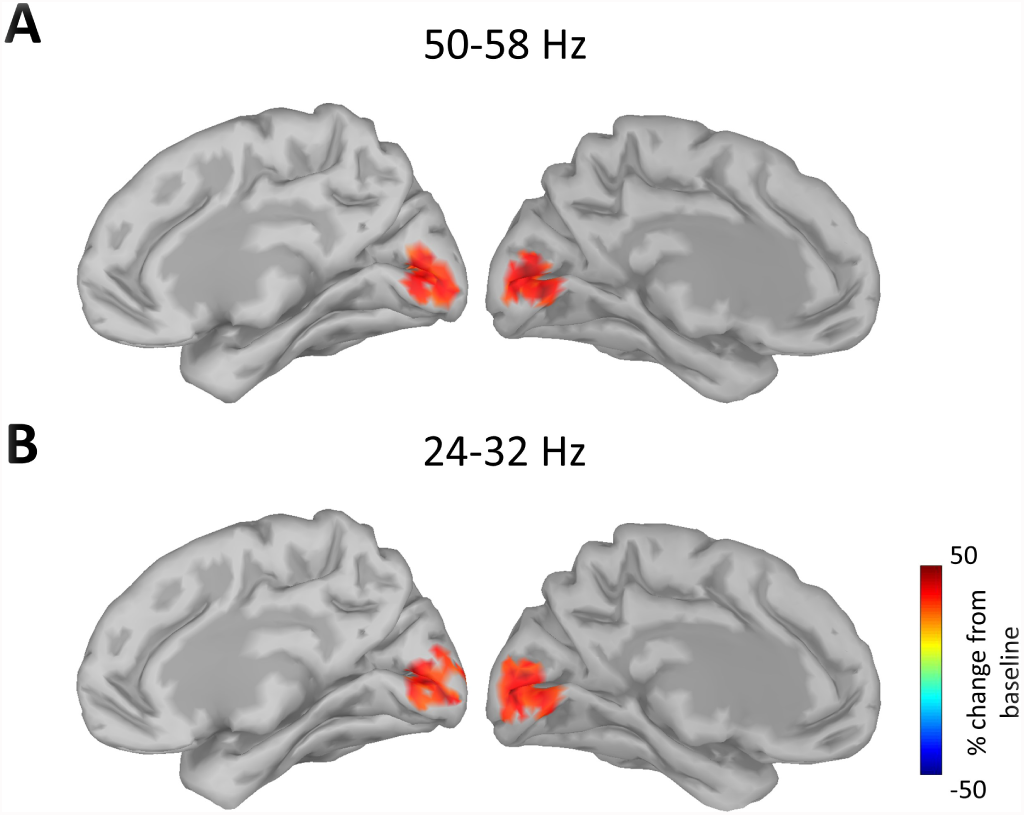
Cortical maps of MEG induced responses at **A)** 50-58Hz and **B)** 24-32Hz. Maps were averaged across the 14 participants, the 6 stimulus conditions, and across time from 300 ms to +800 ms with respect to stimulus onset.

We then conducted region-specific analysis to further investigate stimulus-specific evoked and induced responses in early visual cortex (EVC). Such analysis enabled the study of neuronal activity in EVC directly, though at the cost of averaging over multivariate spatial patterns across the pericalcarine cortex.

EVC induced activity in the 10-80 Hz range is shown in Fig. 7A. Early responses were predominantly contained within the 50-58 Hz band, consistent with our decoding results and prior studies (Adjamian et al., 2004; Muthukumaraswamy and Singh, 2013; Perry et al., 2013). Responses in the 24-32 Hz band were also evident and became progressively stronger towards the end of stimulus presentation. We also observed cardinal orientations having overall weaker induced responses than oblique orientations, termed an inverse oblique effect (Koelewijn et al., 2011).

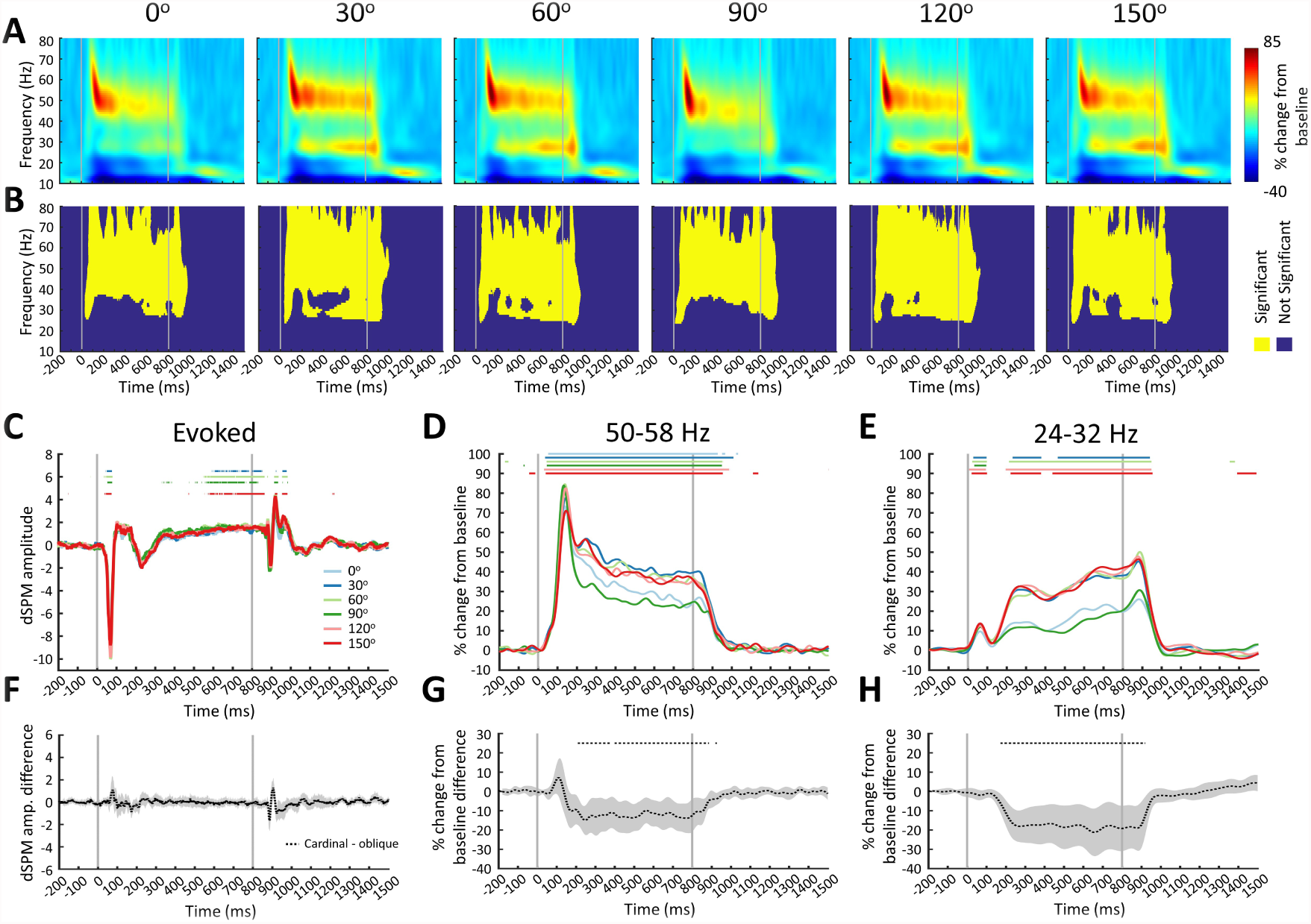
**Fig. 7**. Early visual cortex activity for the 6 Cartesian square-wave gratings. **A)** Time-frequency power maps of induced brain activity in early visual cortex. **B)** Statistical significance maps (*N* = 14; one-sided sign-permutation test; *p* < 0.005 cluster defining threshold; *p* < 0.05 cluster threshold). **CDE)** Evoked response and 50-58 Hz and 24-32 Hz induced response in early visual cortex. **FGH)** Cardinal minus oblique signals in early visual cortex for the corresponding brain responses. (*N* = 14; two-sided sign-permutation test; *p* < 0.05 FDR corrected over time).

Time series of evoked and induced responses for the two frequency bands are shown in Fig. 7CDE. The evoked activity was very similar for all 6 Cartesian grating stimuli, with no consistent preference for cardinal over oblique stimuli (Fig. 7F). The initial 50-58 Hz gamma response around 100 ms after stimulus onset was stronger for cardinal than oblique stimuli (Fig. 7G). While this was not significant, it is consistent with prior findings (Koelewijn et al., 2011). Notwithstanding this transient effect, induced responses in both frequency bands were dominated by an inverse oblique effect for the entire duration of stimulus presentation.

Note that the reverse relation, with cardinal orientations having weaker induced responses (Fig. 7DC) but stronger decoding results (Fig. 3CD) than oblique orientations, is not contradictory. Multivariate analyses are particularly effective in capturing consistent pattern differences and translating them to a positive discrepancy measure, the decoding accuracy. In this case, consistently weaker gamma oscillations for cardinal than oblique orientations resulted in robust decoding results between them.

Taken together, we found a strong inverse oblique effect in induced EVC brain activity, namely weaker responses for cardinal than oblique orientations, but no orientation selectivity in evoked EVC responses. Since MEG signals were summarized to a single time series in EVC, the current analysis also served as a univariate measure of orientation selectivity in EVC. As such, univariate signals had limited sensitivity to orientation information, in contrast to multivariate pattern analyses that were able to pairwise decode all stimuli in the previous sections.

## 5 Discussion

### 5.1 Summary

In this study, we have shown that multivariate methods are well suited for evaluating orientation information. Decoding revealed two frequency bands, 50-58 Hz and 24-32 Hz, with orientation information. Formal comparison using representational similarity analysis with two models of hypothesized orientation representations determined that evoked, and 50-58 Hz and 24-32 Hz induced responses largely shared information, with the cardinal model predominantly explaining their representations. Overall, multivariate methods constituted a principal approach to study orientation information. Such methodological framework can motivate future studies to investigate the encoding of other types of stimulus information to gamma responses.

### 5.2 Multivariate pattern analysis methods are well suited for resolving orientation information in both evoked and induced responses

Previous studies investigating the complex visual processes in the human brain using multivariate pattern analyses methods have focused on decoding visual information from evoked responses, clarifying the representation of objects (Carlson et al., 2013; Cichy et al., 2014; Clarke et al., 2014; Kaneshiro et al., 2015) and scenes (Cichy et al., 2016; Groen et al., 2017), the temporal maintenance of visual information (King et al., 2016), and visual motion (Bekhti et al., 2017). In closer relation to the work described here, MEG evoked responses have been shown to discriminate orientation information of grating stimuli (Ramkumar et al., 2013; Cichy et al., 2015; Wardle et al., 2016).

Here we extended prior findings by demonstrating that multivariate pattern methods are well suited for the analysis not only of evoked but also of induced responses. In fact, we were able to discriminate all pairs of the 6 Cartesian grating stimuli, regardless cardinal or oblique, both from evoked and induced responses, as evidenced by multi-class SVM decoding in Suppl. Figure 4. This highlights the robustness of multivariate pattern analysis in extracting visual information from MEG data.

Our work builds upon a previous study showing that two oblique (+45° and -45°) orientations can be decoded from the sustained gamma response of a source-reconstructed MEG signal in primary visual cortex (Duncan et al., 2010). This study used orientation stimuli that subtended only 1.5° to activate a focal EVC region, and constructed pattern vectors along the spectral dimension using frequency bins in the 20-70Hz band. Our study differed in stimuli and methodology, and aimed to show that gamma patterns in the spatial (and not only spectral) dimension encode stimulus information. We used large (6.5° radius) grating stimuli activating an extended patch of the pericalcarine cortex. Thus, our multivariate classification was sensitive to spatially extended neuronal patterns. Further, we constructed pattern vectors using measurements from the entire MEG sensor array and in some cases spectral values along frequency bands, thus our SVM analyses relied on both spatial and spectral neuronal patterns.

In contrast to the multivariate analysis results, orientation information was very limited in univariate signals from the early visual cortex. We detected no orientation information in evoked EVC responses. The only orientation information in induced EVC responses was an overall stronger gamma power of oblique over cardinal orientations (inverse oblique effect), but otherwise all oblique stimuli had comparable gamma power and were not resolvable from each other. This further highlights that, due to its higher sensitivity, multivariate pattern analyses may reveal encoded information where univariate analyses do not (Haynes, 2015).

A comparable univariate analysis performed directly on sensors corroborated these findings. Sensor topography plots (Suppl. Figure 5) revealed that sensors with the highest induced gamma responses clustered over the occipital cortex. Investigating the 4 sensors with the strongest induced responses affirmed that gamma power was stronger for oblique than cardinal orientations. However, all oblique stimuli had comparable gamma power and were not resolvable from each other.

Since we constructed multivariate pattern vectors from the entire sensor array, in principle the decoded signals could have originated from anywhere in the cortex. However, the origin of visual gamma oscillations is well known to localize in the primary visual cortex (Adjamian et al., 2004; Hoogenboom et al., 2006; Koelewijn et al., 2011), as was confirmed here, too, by source localization (Fig. 6). In agreement with these results, the SVM classifiers were spatially specific to occipital sensors (Suppl. Fig. 6).

We note that in a prior study, using equivalent multivariate analyses procedures and similar grating stimuli, we did not identify cardinal versus oblique effects in MEG evoked responses (Fig. 2 in Cichy et al., 2015). How can this discrepancy be explained? One potential source is differences in the stimulus material. While the stimuli sampled the same orientations as here (0° to 150° in steps of 30°), they differed in spatial frequency, type (sine-wave vs. square-wave), and spatial extent. Presentation time was also considerably shorter (100 ms vs. 800 ms). A probably more critical factor was the difference in signal-to-noise ratio (SNR), with the prior study collecting only 33 trials per stimulus, considerably lower than the 352 trials per stimulus collected here. Further work, investigating the encoding of orientation information from gamma responses as a function of stimulus duration and SNR, might shed light on this issue.

### 5.3 Orientation selectivity of gamma induced responses

RSA enabled the evaluation of two formal models of orientation representation, a categorical cardinal model, and an ordinal angle disparity model. The cardinal model was the dominant model, explaining the MEG 50-58 Hz and 24-32 Hz induced responses very close to noise ceiling (Fig. 5CD). Conversely, the angle disparity model proved a poor model for the induced responses, having very weak correlations with the data. Thus our results indicate that for orientation, a basic and fundamental stimulus feature, the information encoded in MEG induced responses is almost entirely explained by a cardinal versus oblique categorical effect.

This result is consistent with the univariate analyses of EVC induced responses, which found that visual gamma oscillations where weaker in response to cardinal than oblique stimuli (Fig. 6). Termed inverse oblique effect, this finding replicates a prior study (Koelewijn et al., 2011). A possible explanation may be the differential tuning between oblique and cardinal cells. While cardinal orientations may be encoded by a larger cell population (Pettigrew et al., 1968; Maffei and Campbell, 1970), their sharper tuning curves (Rose and Blakemore, 1974) can lead to overall reduced neuronal activity. This is because oblique stimuli will induce firing to a larger assembly of neurons weakly tuned to a broad range of orientations, whereas cardinal stimuli will induce firing to a smaller assembly of neurons specifically tuned to that cardinal orientation.

Even though cardinal orientations dominated the representational structure of the induced response, all pairs of stimuli were decodable. This includes pairs of stimuli with oblique orientations, which were resolved with approximately 55-65% decoding accuracy (Fig. 3CD), despite their EVC gamma power being the same. This highlights the fact that information on orientation is encoded at distinct spatial patterns across the cortex, with measures of overall gamma power and frequency in large patches of cortex failing to capture this information.

The precise nature of the source of oblique versus cardinal effects in the visual cortex remains equivocal. The reasons are multifold, including the presence of different types of perceptual biases (Essock, 1980); the strong dependence of orientation anisotropies to stimuli, with simple stimuli often associated with greater sensitivity to cardinal orientations and natural stimuli to oblique orientations (Essock et al., 2003); the overall weak neural substrate of orientation anisotropies, typically observed in simple cells with small receptive fields (Leventhal and Hirsch, 1980; Orban and Kennedy, 1981); and the disparate methodological approaches (Maloney and Clifford, 2015). Here we have shown that methodological approaches with univariate analysis of induced responses may not sufficiently capture the spatial patterns of orientation selectivity. Thus, we hope to motivate future studies of orientation anisotropies to use multivariate approaches, capable of resolving spatially distributed neuronal patterns. Such investigations would highlight the dependence of orientation information to contrast (Maloney and Clifford, 2015), differentiate information encoded in gamma power versus frequency peak (Jia et al., 2013), and identify whether MEG/EEG can capture broadband frequencies that encode stimulus relevant information (D. Hermes et al., 2015; Ray and Maunsell, 2011), beyond the narrowband frequencies detected here.

### 5.4 Relationship between induced 50-58 Hz and 24-32 Hz bands

Multivariate pattern analysis greatly facilitated the assessment of orientation information in a broad range of frequency bands, with the decoding time-frequency maps (Fig. 2B) revealing a distinct separation of two relevant frequency bands, 50-58 Hz and 24-32 Hz. Substantial evidence confirms they are related to each other, possibly reflecting different aspects of the same underlying neuronal process. First, the 24-32 Hz band was a subharmonic of the 50-58 Hz band, with the ratio of high/low frequency peaks being approximately 2 separately for each subject, despite across-subject variability in frequency peaks (Suppl. Fig. 3). Second, the two bands had overall similar representational structure, with the cardinal model predominantly explaining their representations. Third, the two bands co-localized in EVC (Fig. 6).

While the existence of two orientation-selective bands was unexpected, the result is not original but rather may have been overlooked in literature, with prior studies pointing to similar findings. Hoogenboom et al., (2006) identified two clearly separate visually induced gamma bands in some of their subjects, though at a higher range (one band around 40 Hz and another around 70-80 Hz). Numerous other studies show spectra with two visual gamma bands, though the lower one is typically weak and not discussed (examples include Fig 3A in Hadjipapas et al., 2007; Fig. 1 in Perry et al., 2013; and Fig. 3 in Tallon-Baudry, 2004). Notwithstanding these results, the strength of the 24-32 Hz band in our study is unusual and more investigation will be necessary to identify whether a particular selection of experimental parameters contributed to this effect.

Periodic sensory input, such as tactile vibrations and periodic auditory noise clicks, are known to produce nonlinear responses characterized by components at different harmonic frequencies (Khan et al., 2015; Langdon et al., 2011; Spencer et al., 2008). However, here gamma oscillations were produced by a stationary stimulus and not a periodic sensory input, thus the neural mechanism is probably different. A possible explanation for the generation of both 50-58 Hz and 24-32 Hz rhythms could relate to a progressively reduced firing output of pyramidal excitatory neurons following prolonged periods of visual stimulation, akin to the 800 ms stimulus presentation in our experiment. Over time, pyramidal neurons may not be able to fire fast enough to follow the fast pace of gamma oscillations, and therefore skip cycles leading to a subharmonic. Evidence supporting this view comes from the progressive reduction of 50-58 Hz power over time versus the increase of 24-32 Hz power over time (Fig. 6DE). This reverse behavior could signify neurons initially contributing to the fast rhythm eventually contributing to the slower rhythm.

Given the strong relation between the 50-58 Hz and 24-32 Hz components, we argue that the 24-32 Hz component should be characterized as a lower gamma rhythm, rather than a beta rhythm, which is typically associated with different functional roles.

### 5.5 Multivariate methods offer a principled approach to link gamma responses to the perceptual Gestalt

The precise functional role of visual gamma oscillations remains controversial (Ray and Maunsell, 2015), with studies offering opposing evidence regarding the necessity of gamma oscillations in seeing (Brunet et al., 2014; D. Hermes et al., 2015; Dora Hermes et al., 2015). Here we propose a methodological framework to investigate the role of gamma responses to perceptual Gestalt. We have shown that multivariate spatial patterns of induced gamma responses robustly encoded a simple stimulus feature, the orientation of Cartesian grating stimuli. This opens the possibility of using multivariate methods to study how induced gamma responses could encode other more complex stimulus features.

Such investigations could link the rhythmic firing of neurons in the gamma band to the binding of stimulus features into coherent wholes (Engel et al., 1991; Singer, 1999; Tallon-Baudry and Bertrand, 1999). For example, one could devise models hypothesizing perceptual relations across stimuli, analogously to the cardinal and angle disparity models we devised to test hypothesized relations across orientations. These models could then be compared against the gamma responses, revealing the nature of perceptual information encoded in gamma responses.

### 5.6 Relationship between induced and evoked responses in orientation selectivity

While evoked and induced responses originate from different neural machineries, with evoked components phase-locked to the stimulus and induced responses showing trial-to-trial latency variations, both have been linked to orientation selective neural processing (Arakawa et al., 2000; Koelewijn et al., 2011; Song et al., 2010). In particular, electrophysiological studies have reported event-related potentials with greater amplitude for cardinal than oblique orientations in early P1 and N1 components (Moskowitz and Sokol, 1985; Proverbio et al., 2002), as well as late P2 and P3 components (Yang et al., 2012).

Using multivariate pattern analysis, we extended these studies by obtaining a more sensitive and holistic description of the evoked responses. Our results showed we could discriminate orientation information from evoked responses, with an early transient response the first couple hundred milliseconds after stimulus, followed by more sustained representations later (Fig. 4AD). Orientation information was predominantly explained by the categorical cardinal model, though the ordinal angle disparity model was also significant early in the response.

The functional role of evoked responses, capturing slowly varying neuronal signals, is understood to be different than that of induced responses. Indeed, the angle disparity model dissociated the two responses, indicating that unlike induced responses, early evoked responses differentially encoded orientation with progressively disparate angles. However, for most of the response time, the cardinal model explained both evoked and induced responses near noise ceiling, indicating that both largely encode the same orientation information. Thus, seemingly different responses when viewed from a univariate perspective, provided converging results when studied with a multivariate approach. Future studies, investigating both evoked and induced responses together, may reveal other possible relations in encoding orientation information between the two responses, such as their dependence on stimulus contrast (Maloney and Clifford, 2015).

## 6 Acknowledgements

We are thankful to Krish D. Singh and Suresh D. Muthukumaraswamy for assistance at early stages of this work, and Sheraz Khan and Eric Lowet for discussions on the interpretations of results. This work was funded by the McGovern Institute Neurotechnology Program to D.P. and by the German Research Foundation (DFG, CI241/1-1) to R.M.C. MEG data were collected at the Athinoula A. Martinos Imaging Center at the McGovern Institute for Brain Research, MIT.

